# Floral Cues and Flower Handling Tactics Affect Switching Decisions by Nectar-Foraging Bumble Bees

**DOI:** 10.1101/2024.05.01.592087

**Authors:** Minjung Baek, Jonathan S. Garcia, Daniel R. Papaj

## Abstract

Nectar foraging bees change their use of floral resources as plant species appear in the environment and disappear over their lifetimes. The new flowers used may involve different cues and different nectar extraction tactics. Although bumble bees can adapt to changes in floral cues and required tactics, little is known about whether bees prioritize switching tactics or floral cues when deciding which plant species to switch to. In a laboratory assay, we forced *Bombus impatiens* (common eastern bumble bee) workers either to switch the handling tactic they were using or to continue using the tactic but to switch the colour of artificial flowers foraged on. We examined whether bees’ tendency to change their tactics was influenced by how similar in colour novel flowers were to familiar ones. We conducted a 2 × 2 factorial experiment using artificial flowers, manipulating the handling tactic that bees were initially trained (legitimate visitation or nectar robbing) and the similarity between novel and trained colours (similar or distinct). We found that under most conditions bees preferred to switch flower colours and retain handling tactics. However, when given experience with legitimate visitation and when novel flowers were markedly different in colour from those they had experienced previously, bees tended to switch tactic while continuing to forage on flowers of the same colour. These findings suggest that the similarity in colour of a new floral resource to the currently exploited resource, along with the flower handling tactic employed by bees, jointly plays an important role in decision-making by foraging bumble bees.

## Introduction

Foraging animals frequently face the challenge of the resource environment changing over time and space. Changes include phenological fluctuations in the availability and quality of alternative resources (Cotton, 2007; Lowe et al., 2022; Stephens et al., 2019) and resource depletion by competitors (Balfour et al., 2015; Damas-Moreira et al., 2020; Page & Williams, 2023; Valls-Fox et al., 2018). In such circumstances, foragers may switch from one resource type to another (Lowe et al., 2022; Mitchell, 1990; Page & Williams, 2023). In switching resources, animals must find and recognize new resources, and use them as efficiently as possible. What they learn about finding, recognizing and utilizing their prior resource may affect their choice and manner of utilization of a new resource. There may also be innate biases that affect the pattern of switching.

In this study, we focus on switching of floral resources involved with acquisition of nectar by bees. The availability of nectar varies significantly over time and place (Pleasants & Zimmerman, 1979; Real & Rathcke, 1988; Thomson et al., 1989). What flowers offer nectar will vary over the season as different plant species go through their respective reproductive cycles (Coffey & Breen, 1997; Murphy & Kelly, 2003; Ogilvie & Forrest, 2017; Ogilvie & Thomson, 2016), and as the composition of pollinator fauna exploiting those species changes over time (Balfour et al., 2015; Cane & Payne, 1993; Page & Williams, 2023; Petanidou et al., 2008). Such variation can influence the foraging options available to an individual nectarivore and require them to adapt appropriately (Forster et al., 2023; Heinrich, 1976; Irwin et al., 2010; Lichtenberg et al., 2020; Lowe et al., 2022; Ogilvie & Thomson, 2016; Page & Williams, 2023; Pleasants, 1981).

Bumble bees provide an excellent model in which to examine behavioural aspects of switching food resources. A great deal is known about cues used by bumble bees to find flowers (Chittka, 2022; Rands et al., 2023). How bumble bees perceive colour cues can be assessed using a colour coding system that is widespread in Hymenoptera (Chittka, 1992; Chittka et al., 1992; Lunau et al., 1996). This colour coding system allows us to precisely manipulate the perceived similarity among floral colours by bumble bees. We also know a great deal about the means by which bumble bees extract these nutrients from flowers (Krishna & Keasar, 2019; Laverty, 1994; Laverty & Plowright, 1988; Raine & Chittka, 2007b). Of interest in this study are two distinct tactics, that bees use to extract nectar: legitimate visitation and nectar robbing.

Legitimate visitors use the floral opening to access nectar within the flower while also promoting pollen transfer (Inouye, 1980; Irwin et al., 2010). If a nectarivore enters the same floral opening used by legitimate visitors but does not transfer pollen due to a mismatch of morphologies, it is called nectar theft (Inouye, 1980; Irwin et al., 2010). Alternatively, nectarivores including bumble bees sometimes extract nectar from holes cut in the base of flowers instead of through the main floral opening. Referred to as “nectar robbing”, this behaviour can reduce plant fitness due to lack of pollen transfer (Inouye, 1980; Irwin et al., 2001, 2010; Richman et al., 2021; Rojas-Nossa et al., 2016), but offers benefits to nectarivores by reducing handling time (Dedej & Delaplane, 2005; Lichtenberg et al., 2018) or overcoming physical constraints (Irwin et al., 2010). Nectar robbing can be further categorized into primary nectar robbing and secondary nectar robbing which differ in who is making the holes. Primary nectar robbers make holes by themselves to extract nectar, while secondary nectar robbers use holes made by primary nectar robbers (Inouye, 1980; Irwin et al., 2010).

What guides a bumble bee’s decisions in switching from one plant species to another? Innate and learned responses to floral cues such as colour are important in floral choice by bumble bees (Gumbert, 2000; Lunau et al., 1996; Rodríguez et al., 2004; Schiestl & Johnson, 2013; van der Kooi et al., 2019). Are these responses also important in switching from one plant species to another? Alternatively, are the tactics used to extract nectar from flowers more important in switching to a new species? Do these factors interact in selection of the new resource? We addressed these questions in a laboratory study of *Bombus impatiens*, the common eastern bumble bee. *Bombus impatiens* is a generalist pollinator that forages on a wide variety of floral resources (Russo et al., 2013), and can engage in secondary nectar robbing (Rust, 1979). Bumble bees are known to switch flower types in response to variation in available resources (Cnaani et al., 2006; Heinrich, 1976). However, since switching risks wasting time and energy, bumble bees are also known to stick with familiar options. Such persistence has been termed floral constancy, wherein bumble bees visit the same flower type successively even when equally or more rewarding alternatives are available (Chittka et al., 1999; Grüter & Ratnieks, 2011; Waser, 1986). Just as bumble bees sometimes show floral constancy, they may also show tactic fidelity, in which an individual uses only one flower handling tactic for the majority of visits within a single foraging bout (Bronstein et al., 2017; Lichtenberg et al., 2020).

Previous studies have shown that bees often make foraging decisions based on how certain a floral cue is in predicting the occurrence of floral rewards such as nectar (Leonard et al., 2011; Lynn et al., 2005). Bees might reasonably be expected to determine which novel floral options are worth sampling based on their innate or learned perception of floral cues (Maharaj et al., 2018). We hypothesized that if a cue from a novel flower type was perceived to be a predictor of rewards, bees would continue to use the learned tactic to explore the novel flower type (switch to novel floral cue). Conversely, if a novel cue was not perceived as a predictor of rewards, bees would favor flowers associated with more familiar cues even if they were required to change their handling tactic (switch to the novel handling tactic). To test these hypotheses, we first trained bees to forage on an array that consisted of artificial flowers (hereby referred to as “flowers”) that were rewarding via either legitimate visitation or nectar robbing, but not both. Bees were then exposed to an array of flowers that differed in colour or handling tactic from those on which they had been trained; this new flower array consisted of (a) flowers that were the same colour as those in the original array but required the opposite handling tactic for sucrose reward removal and (b) flowers that were accessible with the same handling tactic as in flowers the original array but were of a novel colour. This manipulation forced individuals either to change the flower type they foraged for or the handling tactic they used in order to obtain a reward. We further manipulated how similar the novel colour was to the original flower colour to change the perceived level of uncertainty of a novel colour in predicting rewards. We predicted that, compared to what bees were trained in the previous phase, bees would be more likely to switch to a novel flower type when the novel colour was relatively similar to the original colour, and more likely to switch to a novel handling tactic on familiar flowers when the novel colour was distinct from the original colour.

## Methods

### Colony Maintenance

We used 68 bumble bee workers from 5 colonies for the experiment. These colonies, purchased from Koppert Biological Systems (Howell, MI, USA) were connected to a network of clear tubing and plywood arenas. Each colony had access to two arenas. The larger arena (82 × 60 × 60 cm) was used for experiments and routine colony feeding; the smaller arena (38 × 60 × 40 cm) was used exclusively for colony feeding, enabling workers to continue foraging while experiments were conducted in the larger arena. The walls of both arenas were painted grey (Glidden, PPG Industries Inc., PA, USA). The setup was illuminated in a 12-hour light, 12-hour dark regimen, using 40 W 4400 lumen LED panels (61 × 61 cm; 5000 K Cool White, James Industry). Colonies had constant access to transparent gravity feeders with 30% (w/w) sucrose solution for regular feeding. We provided 2 to 5 g of ground honey bee pollen (Koppert Biological Systems, Howell, MI, USA) daily in the nest box, with the quantity adjusted roughly in proportion to the brood area (closed cells) of the colony.

### Feeding Array and Artificial Flowers

Throughout the experiment, a 3 × 2 vertical array of automated feeders painted in the same grey colour as the arena was used. The design of feeders was adapted from Kuusela & Lämsä (2016), and consisted of a 3D-printed plastic box (47 × 24 × 35 mm) with a small servo motor inside. The servo motor was wired to a control panel that enabled individual refilling of the feeder with a button press (Figure 1a). When actuated, a motor dipped a loop of wire into a reservoir of 50% (w/w) sucrose solution before raising the loop up through a hole in the top of the feeder. This caused a small volume of sucrose solution (1.74 ± 0.23 µl; mean ± SD; N = 12) to adhere to the wire loop. Artificial flowers were inserted into the hole in the top of the feeder such that the wire loop with sucrose reward was available at the base of the flower (Figure 1b and 1g).

**Figure 1.**
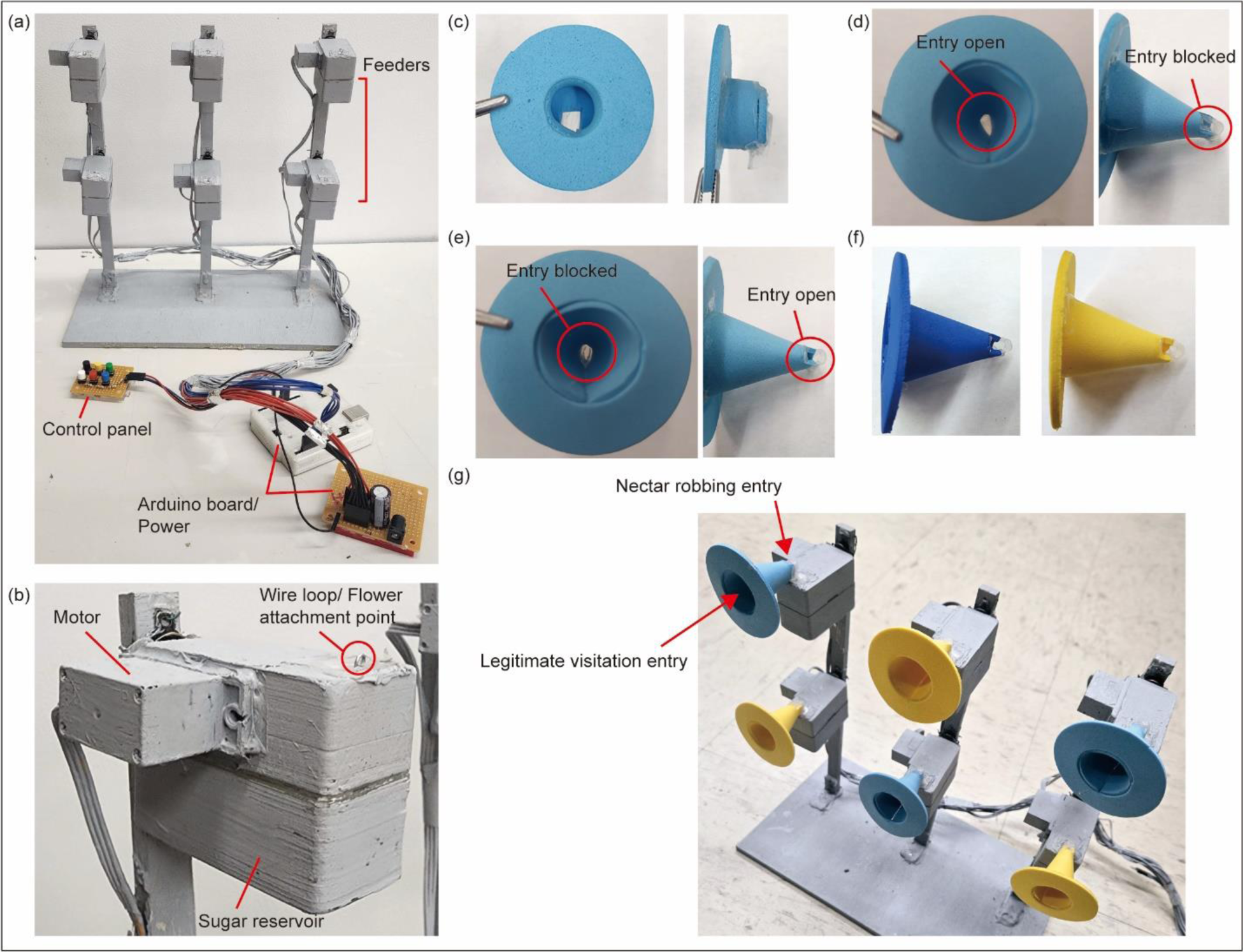
Basic construction of a feeding array, feeders, and artificial flowers. (a) The vertical feeding array consisted of 6 feeders spaced 10 cm apart. Feeders were wired up to an Arduino circuit board controlled by six buttons with each button corresponding to one feeder. (b) Each feeder had a motor connected to a small wire loop and a well holding sucrose solution. By pressing the button, the wire loop was dipped into the sucrose solution, collecting ca. 2 µl of it (1.74 ± 0.23 µl; mean ± SD; N = 12) of it. The loop was then raised back up into the flower on top of the feeder. (c) Pre-training flowers (d) Legitimate flowers and (e) Robbing flowers. Blue flowers are shown here as an example. (f) Novel flowers that differ only in colour compared to training flowers. Dark blue flower and yellow flower represent the similar novel colour (left) and distinct novel colour (right), respectively. (g) Artificial flowers mounted on feeders.

Artificial flowers were constructed from 2 mm foam sheets (Creatology, Michaels Stores, Inc., Texas, USA) and hot glue. Three flower types were constructed: pre-training, legitimate, and robbing flowers (Figure 1c-e; dimensions provided in Figure A1). Pre-training flowers were blue and shaped differently than the other flowers used in the experiment, having a much shorter corolla and no robbing hole. These flowers were designed to train naïve bees to use artificial flowers, while still requiring them to learn to visit a flower legitimately. Both legitimate and robbing flowers had a cone shape with two openings. For legitimate flowers, a small 3 x 4mm opening simulating a robbing hole at the base on the dorsal surface of the flower was blocked, while for robbing flowers, the main opening of the flower was blocked with clear plastic (Figure 1d, e, and g). Sucrose reward could only be accessed via one opening on a given flower, via the wire loop. Both robbing and legitimate flowers were constructed in three colours – blue, dark blue, and yellow (Figure 1d-f). Blue and dark blue constituted similar colours (0.11 hexagon units apart in bee colour space), whereas blue and yellow constituted distinct colours (0.38 hexagon units apart in bee colour space; more details in Figure A2). Flowers were connected to the feeder by a transparent plastic tube, which allowed us to observe whether bees extracted sucrose solution from a wire loop or not.

### Pre-training

To begin pre-training, several naïve bees were introduced to the array with six pre-training flowers by manually lifting them in a vial to the flower opening to forage. After a flower was foraged on and the bee flew away, it was immediately refilled with 50% (w/w) sucrose solution by eye dropper. This process was repeated until bees began to fly to flowers on the array without intervention, at which point a coloured number tag (The Bee Works, Inc., Ontario, Canada) was superglued to the bees’ thorax and they were deemed ready for further training and testing.

### Training and Testing

We employed a 2 × 2 factorial design. One factor was the handling tactic reinforced with sucrose solution in the **training phase**, with bees being trained either to rob blue flowers or to visit blue flowers legitimately. The other factor was the colour pair used in the **test phase**, the blue colour being paired with a novel flower colour. The novel colour was either a similar colour (dark blue) or a distinct colour (yellow) (Figure A2). For either colour pair, a bee trained on a given tactic could only use that tactic to extract sucrose on flowers of the novel colour. To extract sucrose from blue flowers in the test phase, it was required to use the alternative tactic. For example, if the bee was trained to visit blue flowers legitimately in the training phase, it could use that tactic to extract sucrose reward from flowers of the novel colour, say yellow. However, it would be required to rob in the test phase in order to extract sucrose solution from the blue flowers. In total, there were four treatment combinations (Figure 2a): “legitimate trained – similar novel colour” (*n* = 7, 5, 4, and 2 from colony 1, 3, 4, and 5 respectively), “legitimate trained – distinct novel colour” (*n* = 7, 1, 4, 4, and 2 from colony 1, 2, 3, 4, and 5 respectively), “robbing trained – similar novel colour” (*n* = 5, 1, 4, 4, and 2 from colony 1, 2, 3, 4, and 5 respectively), “robbing trained – distinct novel colour” (*n* = 6, 4, 4, and 2 from colony 1, 3, 4, and 5 respectively).

**Figure 2.**
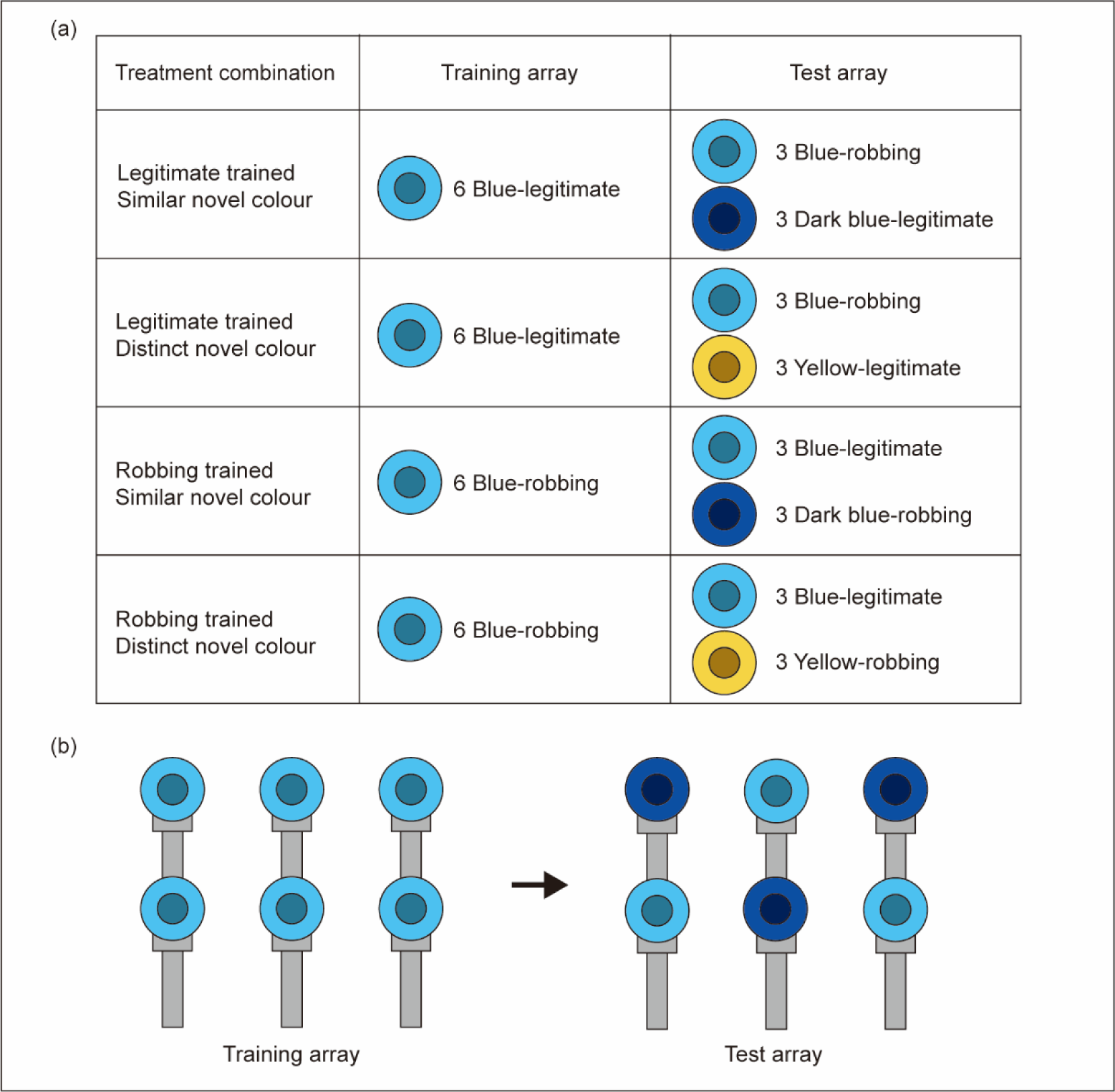
(a) Summary of the four treatment combinations used in the experiment. Each flower had 50% sucrose reward that can be only accessed by either legitimate or robbing tactic. (b) Layout of flowers in the array in training and test phases. This example shows a case where a novel colour is similar to the colour in training (blue). Two flower types were alternated in the test array as shown.

Focal bees were haphazardly assigned to a given training-test combination. Details of the training phase and test phase are given below.

#### Training phase

The large arena was emptied of all bees and pre-training flowers were replaced with six identically constructed blue flowers that could be visited legitimately (hereafter “legitimate” flowers) or robbed (hereafter “robbing” flowers). Each flower offered 50% sucrose solution, and each could be independently replenished with a push of the button assigned to it (see Figure 1). The focal bee was held in the tubing network until the flower array was set up, then released into the arena and allowed to forage freely. Flowers were refilled after a subject visited the flower, extracted sucrose reward, and flew from the flower. Bees that did not extract a reward for more than 20 minutes were removed from the arena and excluded from further assay. If a bee reached more than 25 successful visits in a single foraging bout (*n* = 66), it was advanced to the test phase upon its next return to the arena from the colony; if a bee returned to the colony before 25 visits were reached (*n* = 2), it was allowed back into the arena to continue foraging in the training array until it reached at least 25 visits.

#### Test phase

Bees that completed more than 25 successful visits were allowed back into the colony and held in the tubing network while training-phase flowers were swapped for test-phase flowers. The array was set up with three flowers of the same blue colour used in training but of the opposite handling type and three novel-colour flowers of either similar (dark blue) or distinct (yellow) colour. Access to the 50% sucrose reward from the novel-colour flowers required the same handling tactic adopted in the training phase; access to sucrose from the blue flowers required the other handling tactic. We alternated the two flower types throughout the array as shown in Figure 2b. Flower placement was also alternated over successive trials such that half of the bees in each treatment group foraged on arrays with blue flowers in the first position in the upper row and half foraged on arrays with flowers of novel colour first in that position.

A focal bee was then released into the arena and allowed to forage freely. Flowers were refilled after a subject visited the flower, extracted sucrose reward, and flew from the flower. We filmed the entire session of the test phase using a digital camcorder (Canon VIXIA HF R400). The handling tactic and colour of flower visited were recorded for each visit. If bees tried both tactics on the same flowers, the tactics were sequentially recorded as if they were on different flowers. The test phase continued until the bee attempted to return to the colony or became inactive for at least five minutes. After test, bees were removed from the colony and euthanized.

### Behavioural coding

We recorded legitimate and robbing attempts of bees in the test phase. A legitimate attempt was recorded when a bee fully entered the flower opening (all 6 legs were inside the cone). A bee landing on the outside of the front of the flower but not entering the opening or backing out before fully entering the opening was not counted as a legitimate attempt. A robbing attempt was recorded when a bee landed at the base of the flower near or on the opening or clear plastic cover and probed for an opening to forage from.

Attempts to use these tactics at a given flower were further categorized according to whether bees switched flower colour, switched the handling tactic, or showed no switching compared to what they had been trained to in the training phase. Our focus was on identifying switching from the options available during the training phase, rather than switching among the options provided during the test phase itself. If a bee foraged on a flower of novel colour in the test phase using the same tactic adopted in the training phase, that attempt was categorized as “switching flower colour only.” Foraging attempts on the same colour learned in the training phase but using the different tactic were categorized as “switching handling tactic only.” Foraging attempts on the novel flower colour using the tactic not learned in the training phase were categorized as “switching both flower colour and handling tactic.” Finally, if a bee trained on a given tactic on blue flowers attempted to perform the same handling tactic on a blue flower in the test phase, that visit was categorized as “not switching.” Of these four types of foraging attempts, only two were rewarding: “switching flower colour only” and “switching handling tactic only.”

### Statistical Analyses

We analyzed foraging attempts that bees made in the test phase to determine whether bees switched from what they had been trained in the training phase. First, we compared the observed proportion of bees attempting a trained tactic on flowers with a trained colour on their first attempt to the expected proportion of 25% via the binomial test.

Furthermore, we ran a generalized linear mixed model with a binomial distribution to determine if bees were more likely to switch over the sequence of attempts. The response variable was binary, representing the presence (coded as 1) or absence (coded as 0) of “not switching.” Absence of “not switching” includes the following types of switching attempts: “switching flower colour only”, “switching handling tactic only”, and “switching both flower colour and handling tactic”. The trained-tactic treatment, novel-flower-colour treatment, position of attempt in sequence, and all interactions between these variables were included as fixed effects. Bee identity and colony identity were included as random effects, where Bee identity was nested within colony identity.

To determine how the trained-tactic and novel-flower-colour treatments affect the tendency of a bee to either switch handling tactic or switch flower colour, we ran GLMM with a beta binomial distribution. The ratio of “switch flower colour only” to the sum of occurrences of “switch flower colour only” and “switch handling tactic only” was a response variable. Trained-tactic treatment, novel-flower-colour treatment, and their interaction were fixed effects. Colony identity was included as a random effect.

Lastly, we compared the frequency of “switching both flower colour and handling tactic” over total attempts among the 4 treatment combinations by using GLMM with a beta binomial distribution. Trained-tactic treatment, novel-flower-colour treatment, and their interaction were fixed effects. Colony identity was included as a random effect.

All analyses were completed using R v.4.3.1 (R Core Team, 2023). All GLMMs were conducted using glmmTMB package (Brooks et al., 2017). We used car package to calculate Type 3 sums of squares ANOVA for each term in models (Fox & Weisberg, 2018). After constructing a model, Tukey’s HSD post-hoc comparisons were run using emmeans package (Lenth, 2022).

### Ethical Note

This research adheres to all ASAB/ABS Guidelines for the Use of Animals in Research, the legal requirements of the U.S., and all institutional guidelines. Bees were humanely euthanized by freezing at the end of the experiment.

## Results

### Do bees switch flower colour, handling tactic, both, or neither?

When bumble bees were first introduced to the test array, they attempted to visit the same flower colour and use the same handling tactic learned in the training phase more than the 25% attempts expected by chance (Binomial test against 25% null expectation: *P* < 0.0001; Figure 3). Over the course of their foraging attempts, this behavioural pattern decreased in frequency (GLMM: *estimate ± SE* = 0.02 *±* 0.003, *Z* = – 6.1, *P* < 0.0001; Figure 3). The decrease in frequency was highly variable. One individual attempted their original handling tactic on blue flowers 51 times before attempting a novel tactic, while other individuals engaged in a new tactic or visited the novel colour on their first visit. Bees that had been trained to visit legitimately and were then exposed to yellow, the more distinct novel colour, were more likely to exhibit the previously trained tactic on the previously trained colour at the beginning of the test phase; however, they were also faster in switching to alternatives than were bees in other treatment combinations (emtrends pairwise comparison: all P < 0.0001; Figure 3).

**Figure 3.**
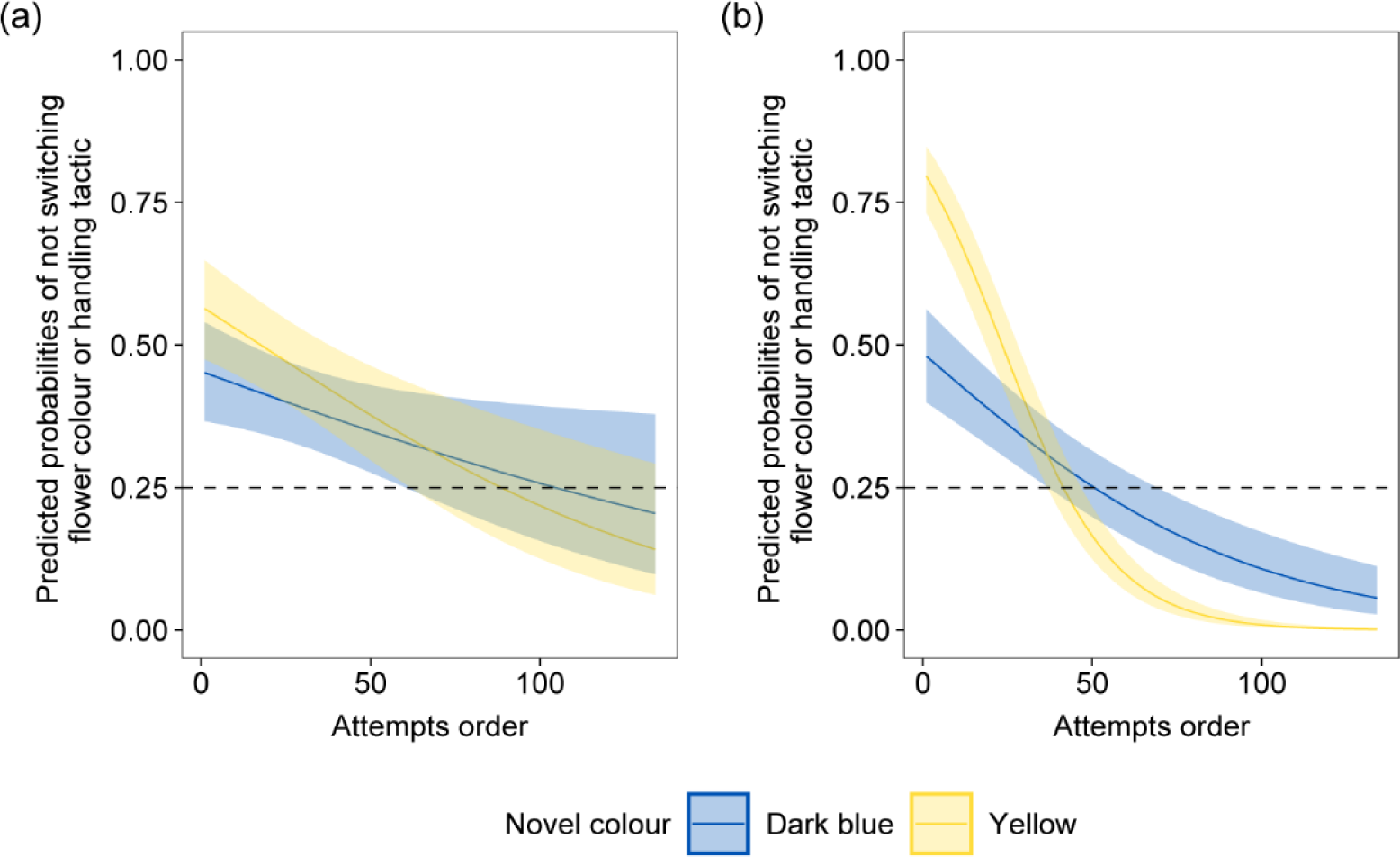
Estimated probabilities of bees not switching flower colour or flower handling tactic over attempts in the test phase. Each prediction was derived from the results of its respective treatment combination. Bees were previously trained to (a) nectar robbing tactic or (b) legitimate visitation tactic on blue flowers in the training phase. Novel flower colours introduced to bees during the test phase were similar to (dark blue) or distinct from (yellow) the trained colour. "Not switching” consisted of doing what was rewarding in the training phase but was no longer rewarding. “Switching” consisted of three types of attempts: two of these (“switching flower colour only” and “switching handling tactic only”) were rewarded and one (“switching both flower colour and handling tactic”) was not rewarded. Lines indicate estimated means and shaded regions indicate 95% confidence intervals. Dashed lines at 0.25 indicate probabilities of “not switching” expected by chance. *N* = 16 for robbing-similar bees, *N* = 16 for robbing-distinct bees, *N* = 18 for legitimate-similar bees, *N* = 18 for legitimate-distinct bees.

### Switch flower colour only vs. switch handling tactic only

With respect to the two rewarding options in the test phase, “switching flower colour only” and “switching handling tactic only”, an asymmetric pattern emerged across treatment combinations. In three of the four treatment combinations, individuals tended to switch flower colour while retaining handling tactic (Figure 4, Table A1). However, bees previously trained to legitimate visitation and given yellow (distinct from trained colour) as the novel colour tended to show fidelity to blue and switch handling tactic (Figure 4, Table A1). The differences are statistically significant (Post-hoc comparison: vs. robbing-similar: *P* = 0.008, vs. robbing-distinct: *P* = 0.014, vs. legitimate-similar: *P* = 0.011; Figure 4, Table A2).

**Figure 4.**
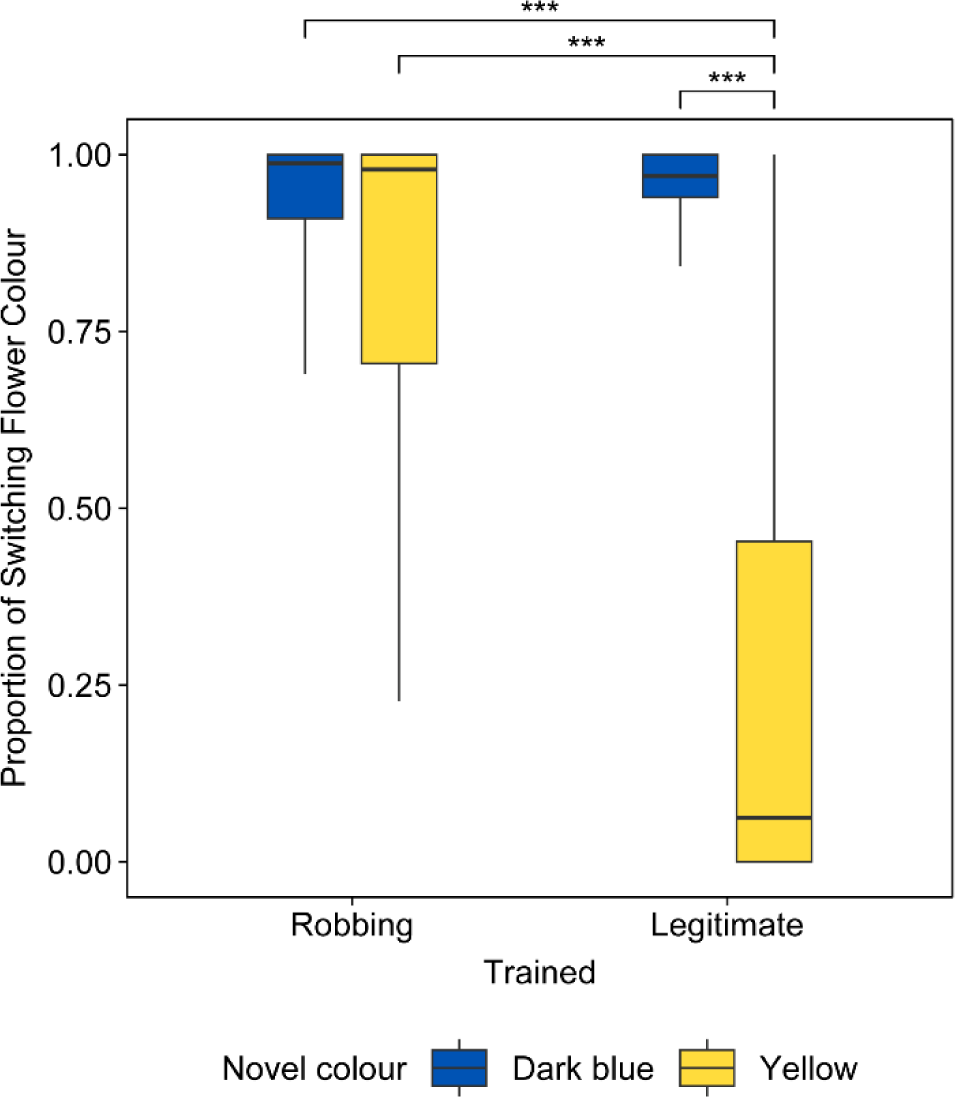
Proportion of rewarding switches that involved switching flower colour only (= (# switching flower colour only)/(# switching flower colour only + # switching handling tactic only)). Bees were either trained to nectar robbing or legitimate visitation tactic on blue flowers in the training phase. Each bee in the test phase was presented with novel flower colours that were either similar to (dark blue) or distinct from (yellow) the trained blue colour. Boxes indicate medians and 25% and 75% percentiles. Whiskers indicate the maximal and minimal data ranges. *N* = 16 for robbing-similar bees, *N* = 16 for robbing-distinct bees, *N* = 18 for legitimate-similar bees, *N* = 18 for legitimate-distinct bees. *** indicates p-value < 0.001 (Tukey’s post hoc test).

### Switch both flower colour and handling tactic

Switching both flower colour and switching handling tactic from what bees had learned during the training phase was the least common of the four possible patterns of switching (Figure A3). This unrewarding type of attempt occurred once or more in 40 out of 68 tested bees (Figure 5). Occurrences were unevenly distributed among treatment combinations. This behaviour occurred in 17 out of 18 bees trained in legitimate visitation and presented with distinct novel colours, but occurred in 56% or less of bees in the other combinations (25% in robbing-trained/similar-colour, 56% in robbing-trained/distinct-colour, 56% in legitimate-trained/similar-colour). In legitimate-trained/distinct-colour bees, this behaviour usually appeared after a switch in handling tactic (Table A3). Also, bees in this treatment combination made this attempt more overall compared to other combinations (Post-hoc comparison: vs. robbing-similar: *P* < 0.0001, vs. robbing-distinct: *P* = 0.0006, vs. legitimate-similar: *P* = 0.0002; Figure 5, Table A4)

**Figure 5.**
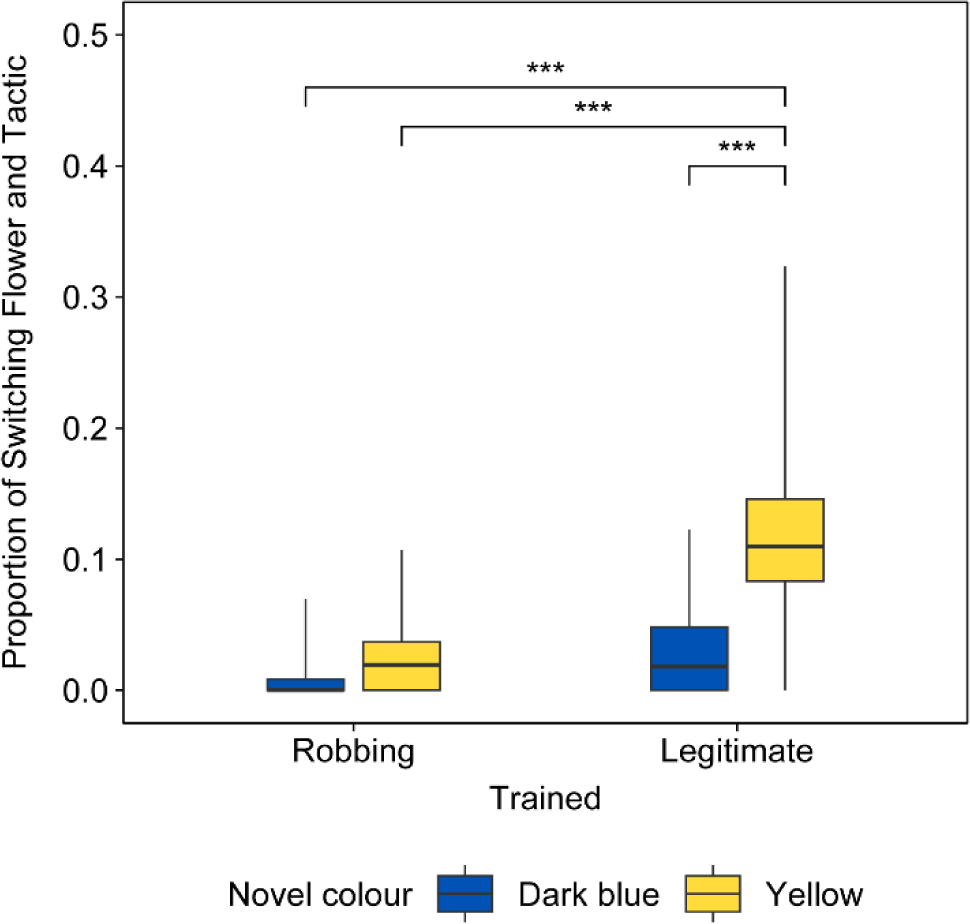
Proportion of switches that involved switching both flower colour and handling tactic (= (# switching both flower colour and tactic)/(# total attempts both rewarding and unrewarding)). Switching both flower colour and handling tactic was unrewarding. Bees were either trained to nectar robbing or legitimate visitation tactic on blue flowers in the training phase. Boxes indicate medians and 25% and 75% percentiles. Whiskers indicate the maximal and minimal data ranges. *** indicates significant differences at p = 0.001 level according to a Tukey’s HSD test. *N* = 16 for robbing-similar bees, *N* = 16 for robbing-distinct bees, *N* = 18 for legitimate-similar bees, *N* = 18 for legitimate-distinct bees

## Discussion

### Fidelity to and Switching of Floral Resources

Multiple hypotheses have been proposed to explain the observed consistency of bees in visiting the same type of flowers and employing the same flower handling tactic (Bronstein et al., 2017; Chittka et al., 1999; Grüter & Ratnieks, 2011). One such hypothesis suggests that being consistent in one option reduces costs associated with exploration and learning (Chittka et al., 1999; Grüter & Ratnieks, 2011). Nevertheless, floral resources in nature are ephemeral and bees do transition from one plant species to another (Heinrich, 1979; Ishii & Kadoya, 2016; Ogilvie & Thomson, 2016; Raine & Chittka, 2007a), giving incentive to understand the cognitive processes involved in switching from one type of flower to another. To make that assessment, we forced bees in the test phase to switch handling tactic or flower colour in order to receive a sucrose reward. Fidelity to both floral type and handling tactic was not a rewarding option. We found that bumble bees were generally successful in switching either tactic or flower colour when the original tactic/colour combination was no longer rewarding. Whether they switched and how they did so depended on the tactic they had been first using and the visual resemblance of novel flowers to familiar flowers.

### Generalization and Innate Biases

In line with our prediction, bees overall tended to retain their trained tactic and switch immediately to novel flowers whose colour resembled that of previously experienced flowers, regardless of the trained tactic. This behaviour suggests that bees generalized the colour of the original blue flowers to the dark blue flowers, even though those two colours are believed to be perceptually discernible to bees (Dyer & Chittka, 2004). Stimulus generalization is a universal phenomenon in animals (Ghirlanda & Enquist, 2003), including in bees’ responses to floral stimuli (de Premorel et al., 2017; Gumbert, 2000; Lynn et al., 2005; Muth et al., 2015; Pietrantuono et al., 2019). Our results are in line with those of Chittka et al. (1997) who observed that bees were more likely to transit between flower species that were visually similar.

However, generalizing stimuli incurs a risk of making incorrect responses (Darst & Cummings, 2006). Different stimuli might indicate a need to make distinct responses, and in such cases, generalization is not desirable. Thus, we predicted that distinct differences between stimuli would deter bees from generalizing, and cause bees to be faithful to their original colour type. Our result was consistent with this prediction in the case of bees trained to visit flowers legitimately. Rather than adapting to the novel colour, bees continued to forage on the original blue flowers by employing a novel tactic. Other studies have also demonstrated that bumble bees are able to adjust their flower handling tactics as required, often involving switching to a newly-rewarding tactic (Baek et al., 2023; Lichtenberg et al., 2020). These results indicate that bees might perceive novel floral cues as different from what they previously experienced and therefore not necessarily predictive of rewards, causing them to focus their foraging efforts on familiar flowers.

Contrary to our prediction, we observed that nectar robbers were more likely to switch flower colours than switch to legitimate visitation on the same flower colour, even when the colour of the novel flowers was markedly different from that of the trained flowers. This result suggests that bees switch more readily from legitimate visitation to nectar robbing, than from nectar robbing to legitimate visitation. The asymmetric nature of tactic switching could also explain the high frequency of “switching both flower colour and tactic” behaviour observed in bees trained to the legitimate tactic and presented with distinctly coloured novel flowers (Figure 5). As legitimate trained bees initially switched their tactic on the trained flowers, this newly acquired robbing tactic could be readily transferred to flowers with novel colours, resulting in these bees switching both flower colour and tactic. This transition did not occur with robbing-trained bees, as a switch from robbing to legitimate visitation was rarely observed after they switched flower colours.

Why would nectar robbers be more likely to switch to the novel flower colour than switch to the novel tactic? One relevant aspect of our experimental apparatus was our use of grey colours to paint feeders on which artificial flowers were mounted. Nectar robbers in our study may have used the grey colour of the feeders as a cue to locate flowers during the training phase, and continued to rely on this cue in the test phase as well. This response to feeder cue might have facilitated switching to novel flower colours while maintaining nectar robbing tactic.

Importantly, such circumstances are pertinent to what pollinators encounter in nature. Flowers are often robbed at the base of the corolla, near the sepals and pedicels. The sepals and pedicels of many species have a similar green colour and form, possibly providing a visual cue that nectar robbers respond to, but which legitimate visitors may not.

Another possible factor in our asymmetric results is the occurrence of innate biases of continuing specific motor routines in bees. Bumble bees and honey bees are well-studied for their innate preferences for specific floral traits (Giurfa et al., 1995; Gumbert, 2000; Howard et al., 2019; Lehrer & Campan, 2005; Lunau, 1992; Lunau et al., 1996; Rodríguez et al., 2004) and for innate dispositions in the learning of tactics for flower handling (Laverty & Plowright, 1988; Russell et al., 2016). We know less about innate dispositions in the switching of preferences and tactics. Perhaps the nectar robbing tactic is inherently prone to be transferred across different flower types, compared to legitimate visitation. During our training phase, some naïve bees were slow to use cone-shaped artificial flowers legitimately. They often terminated a legitimate visit before the reward was reached, crawling back from the entrance. This behavior disappeared once bees successfully obtained a sucrose reward via legitimate visitation. Robbing-trained bees might have been similarly deterred from visiting artificial flowers legitimately. Although possible, we believe such deterrence had a minimal effect on our results because robbing-trained bees typically did not land on the side of the flower where the legitimate entrance was located and thus were unlikely to attempt to enter the entrance in the first place.

### Future Directions: Social Information, Reward Value, and Costs of Learning

Our findings suggest several future directions. Future research could examine how social information influences the tendency to switch flower type versus extraction tactics. In our study, bees were tested independently and did not interact with other foragers. However, avoidance of novel flowers has been shown to be mitigated by the presence of conspecific bees (Kawaguchi et al., 2007), and novel tactics have been shown to be acquired through observation of conspecific behaviour (Leadbeater & Chittka, 2008; Loukola et al., 2017). Foraging bees might also influence other bees by bringing floral odors back to the colony (Dornhaus & Chittka, 1999).

Another future direction would be analysis of reward value in relation to switching. In environments where resources change in availability over time and space, animals are expected to adjust their foraging strategy to increase their net gains. Given that pollinators adjust their behaviour according to the quality and quantity of rewards they obtain (Cnaani et al., 2006; van der Kooi et al., 2021), it is possible that variation in reward value might affect the decision in switching observed in this study. In our experiment, bees altered their flower or tactic choices in response to abrupt changes in rewards, and both transitions resulted in sucrose rewards of the same volume and concentration. However, the value of rewards associated with each choice might not be the same in the environment that bumble bees face. For example, if flowers that bees were foraging on have been completely depleted, switching to a flower of novel colour could be favored, as switching to a novel tactic on the original flowers would no longer yield any rewards. Conversely, if the novel flower species either lacks rewards or offers less rewards than the original species and alternative tactics provide a way to extract rewards from the original species, then switching tactics could be preferred.

It is worth noting that our experiment exclusively considered nectar as reward. However, pollinators often obtain other type of resources from flowers as well (e.g., pollen, heat, odour, and oil; Simpson & Neff, 1981) and the need to collecting alternative resources might affect the decision to switch tactics. For example, bees that collect both pollen and nectar might prefer switching from nectar robbing to legitimate visitation, as nectar robbing tactic bypasses contact with pollen.

In addition to reward value, costs associated with learning new flower handling tactics might affect the tendency of bees to switch. Our study was restricted to two tactics, legitimate visitation and nectar robbing on simple cone-shaped flowers. Yet the motor routines required to extract rewards from flowers vary depending on flower morphology. Visiting one flower type may require different motor routines than another, even if both routines fall under the same category of "legitimate visitation" or "nectar robbing" (Ishii & Kadoya, 2016; Krishna & Keasar, 2019; Laverty, 1994). Becoming efficient in these motor routines, especially for flowers with complex morphologies, could incur a significant cost of time and energy for pollinators (Krishna & Keasar, 2019; Laverty, 1994), deterring them from switching to a novel motor routine.

Furthermore, similarities in motor routines in co-flowering species, alongside the inherent complexity of each motor routine, might also affect bees’ decisions to switch, as bees can transfer motor routines they have learned from one species to another, reducing the cost of learning (Ishii & Kadoya, 2016; Krishna & Keasar, 2021; Richman et al., 2021).

### Conclusion

Bees and other pollinators forage in a highly diverse ‘biological market’ of flowers that is changing over time and space. As rewarding species disappear and new ones appear, and the composition of the pollinator community changes, foragers must adjust their behaviour. They may need to attend to new floral signals and/or to utilize new means to extract nectar or pollen. Previous studies have separately investigated whether bumble bees can change their flower handling tactics or their responses to floral signals. To our knowledge, this is the first study to examine variation in floral signals and handling tactic together. Not only do changes in floral signals and flower handling requirements each contribute to switching by bees, but their effects interact. The form of the interaction in our study was complex, only partly meeting *a priori* predictions, which underscores the value of investigating such interactions. Our findings propose a novel perspective regarding how the composition of neighboring flowers, combined with initial experience of pollinators, may shape decision making by these pollinators.

## Data availability

Data and R code associated with this study are available on OSF (https://osf.io/p3zca/?view_only=53da92b76b5f40b5b673f19527ca557f)

## Appendix

**Figure A1.**
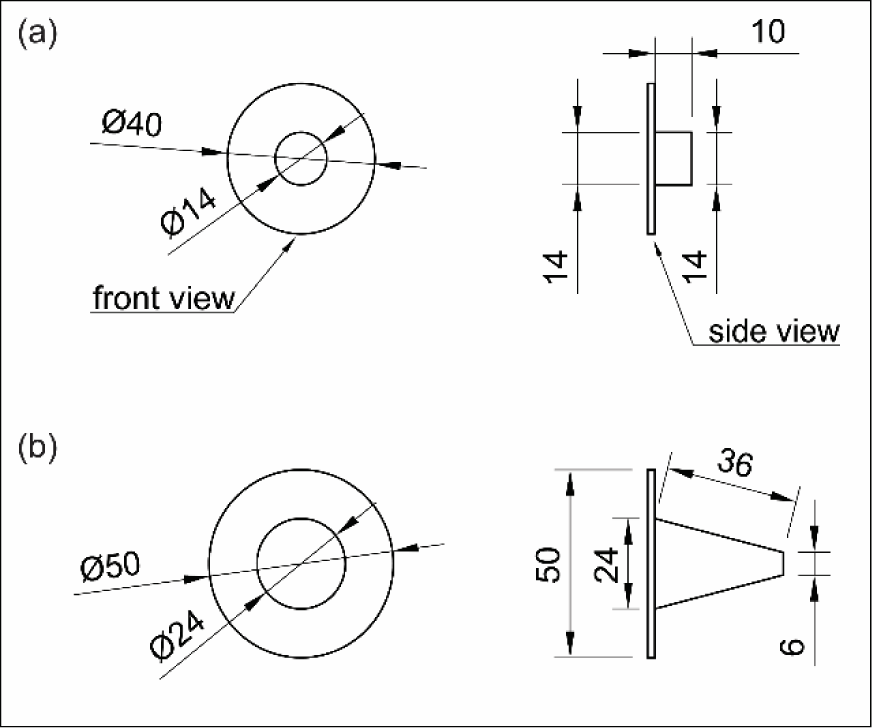
Dimensions of artificial flowers used in the experiment. (a) pre-training flower (b) robbing and legitimate flowers.

### Reflectance spectra and bee colour space

We measured reflectance spectra using an UV-VIS spectrophotometer (Ocean Optics USB2000) with deuterium-halogen light source (Ocean Optics DH2000) and 3 ms integration time. The foam sheets used to construct artificial flowers and the wall of the foraging arena were measured. Prior to measurement, calibration of the spectrophotometer was done with a white Spectralon reflectance standard (USRS-99-010 AS-01158-060; LabSphere). To convert the measured reflectance spectra into the bee colour space, we used the pavo package in R (Maia et al., 2013). We constructed the colour space using the colour opponent model (Chittka, 1992), spectral sensitivities for *Bombus impatiens* (Skorupski & Chittka, 2010), and irradiance in the foraging arena. The colour of the foraging arena wall was used as the background colour. Euclidean distances between colours were calculated to quantify perceptual colour differences in bumble bees.

**Figure A2.**
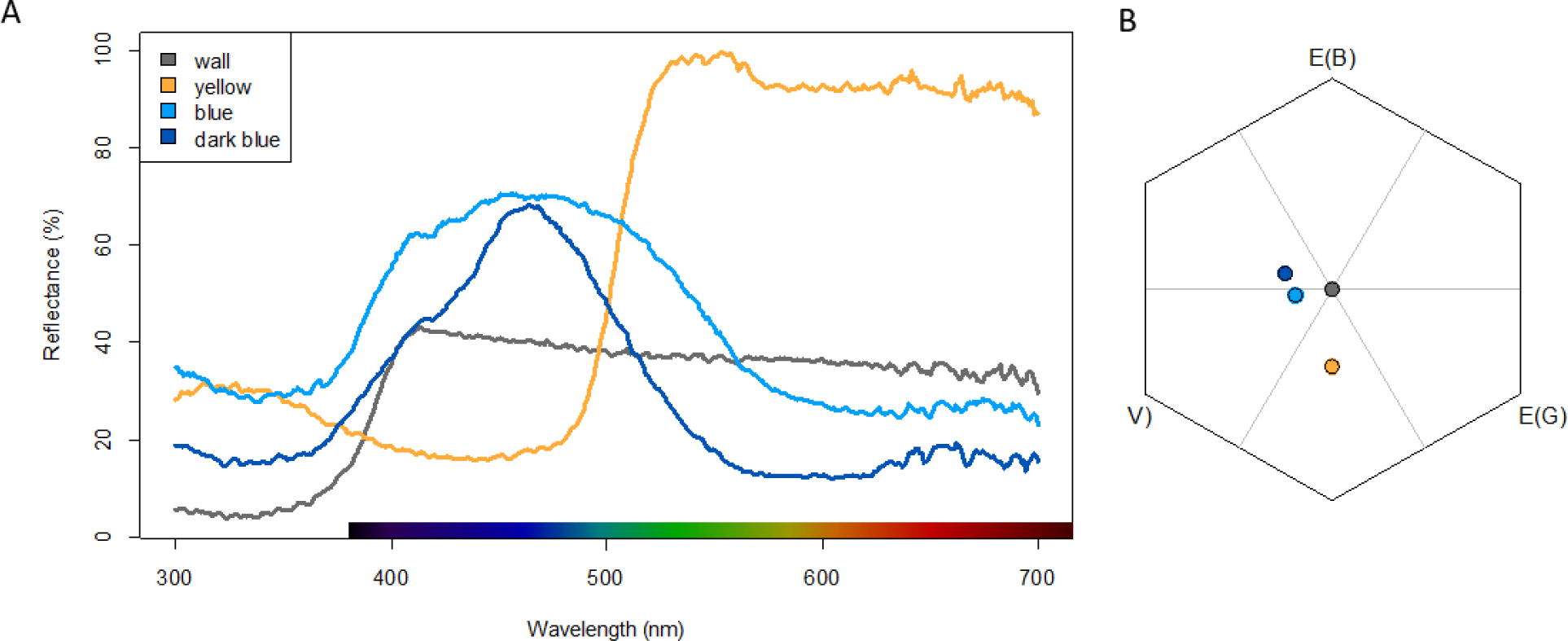
(a) The reflectance spectra and (b) bee colour space of artificial flower colours used in the experiment. “Wall” represents the wall colour of the foraging arena. For reference, a human-based colour spectrum is shown as a bar on the wavelength axis.

**Table A1.** Estimated marginal means on the proportion of rewarding switches that involved switching flower colour only (= (# switching flower colour only)/(# switching flower colour only + # switching handling tactic only)) for each treatment combinations.

| Treatment combination | estimate | SE | Lower 95% CL | Upper 95% CL |
| --- | --- | --- | --- | --- |
| legit-similar | 1.836 | 0.395 | 1.062 | 2.610 |
| legit-distinct | -1.192 | 0.382 | -1.941 | -0.444 |
| rob-similar | 2.033 | 0.437 | 1.177 | 2.890 |
| rob-distinct | 1.612 | 0.430 | 0.770 | 2.454 |

Estimates are given in log-odd ratio. Estimate larger than 0 indicates that bees are more likely to switch flower colour only and estimate smaller than 0 indicates that bees are more likely to switch handling tactic only. CL: confidence limits.

**Table A2.**
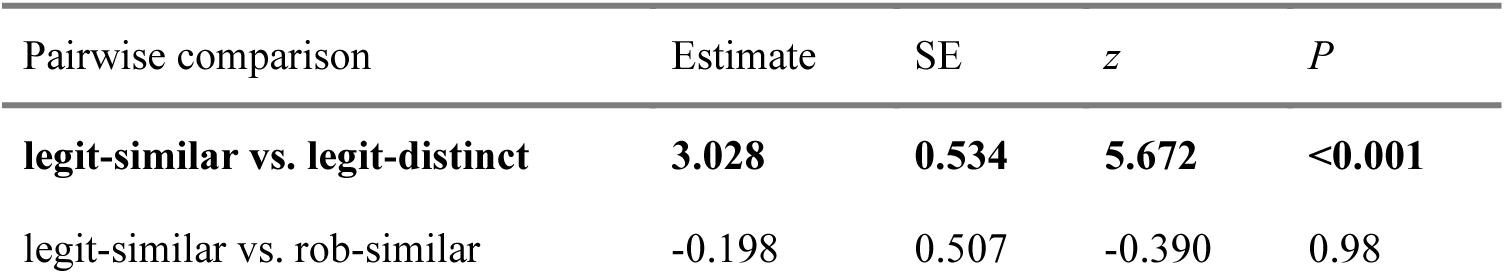
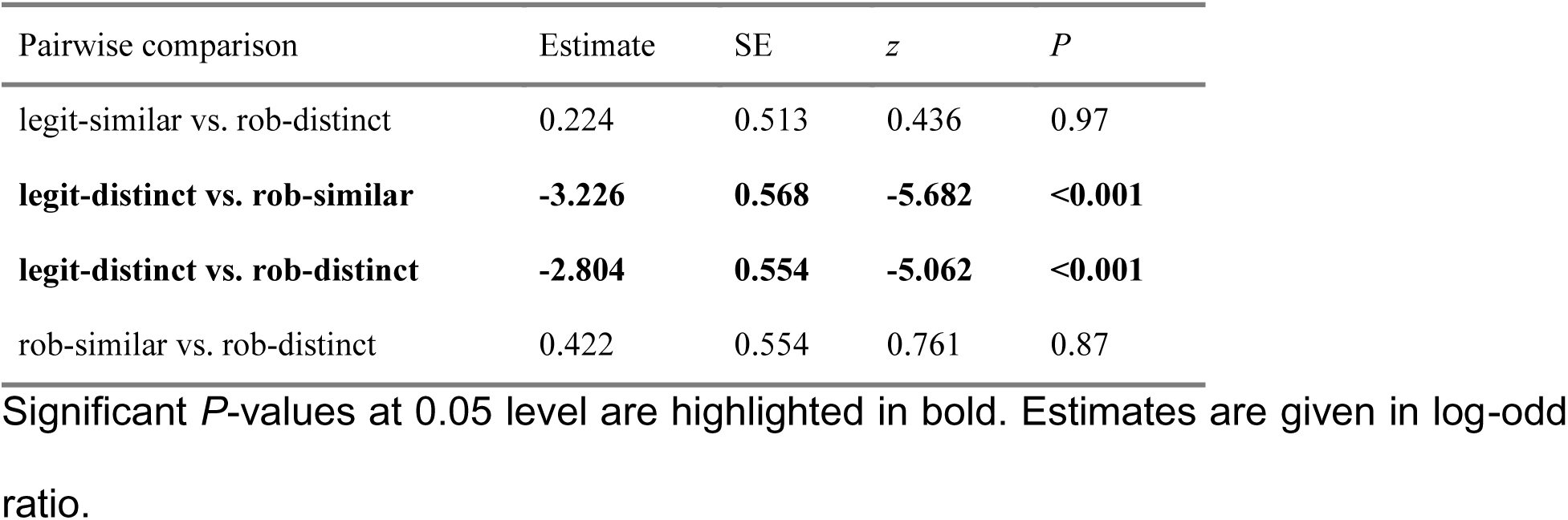
Pairwise comparisons with Tukey HSD adjustment on the proportion of switches that involved switching both flower colour and handling tactic (= (# switching both flower colour and tactic)/(# total attempts both rewarding and unrewarding)) among treatment combinations.

| Pairwise comparison | Estimate | SE | *z* | *P* |
| --- | --- | --- | --- | --- |
| **legit-similar vs. legit-distinct** | **3.028** | **0.534** | **5.672** | **<0.001** |
| legit-similar vs. rob-similar | -0.198 | 0.507 | -0.390 | 0.98 |
| legit-similar vs. rob-distinct | 0.224 | 0.513 | 0.436 | 0.97 |
| **legit-distinct vs. rob-similar** | **-3.226** | **0.568** | **-5.682** | **<0.001** |
| **legit-distinct vs. rob-distinct** | **-2.804** | **0.554** | **-5.062** | **<0.001** |
| rob-similar vs. rob-distinct | 0.422 | 0.554 | 0.761 | 0.87 |

Significant *P*-values at 0.05 level are highlighted in bold. Estimates are given in log-odd ratio.

**Table A3.**
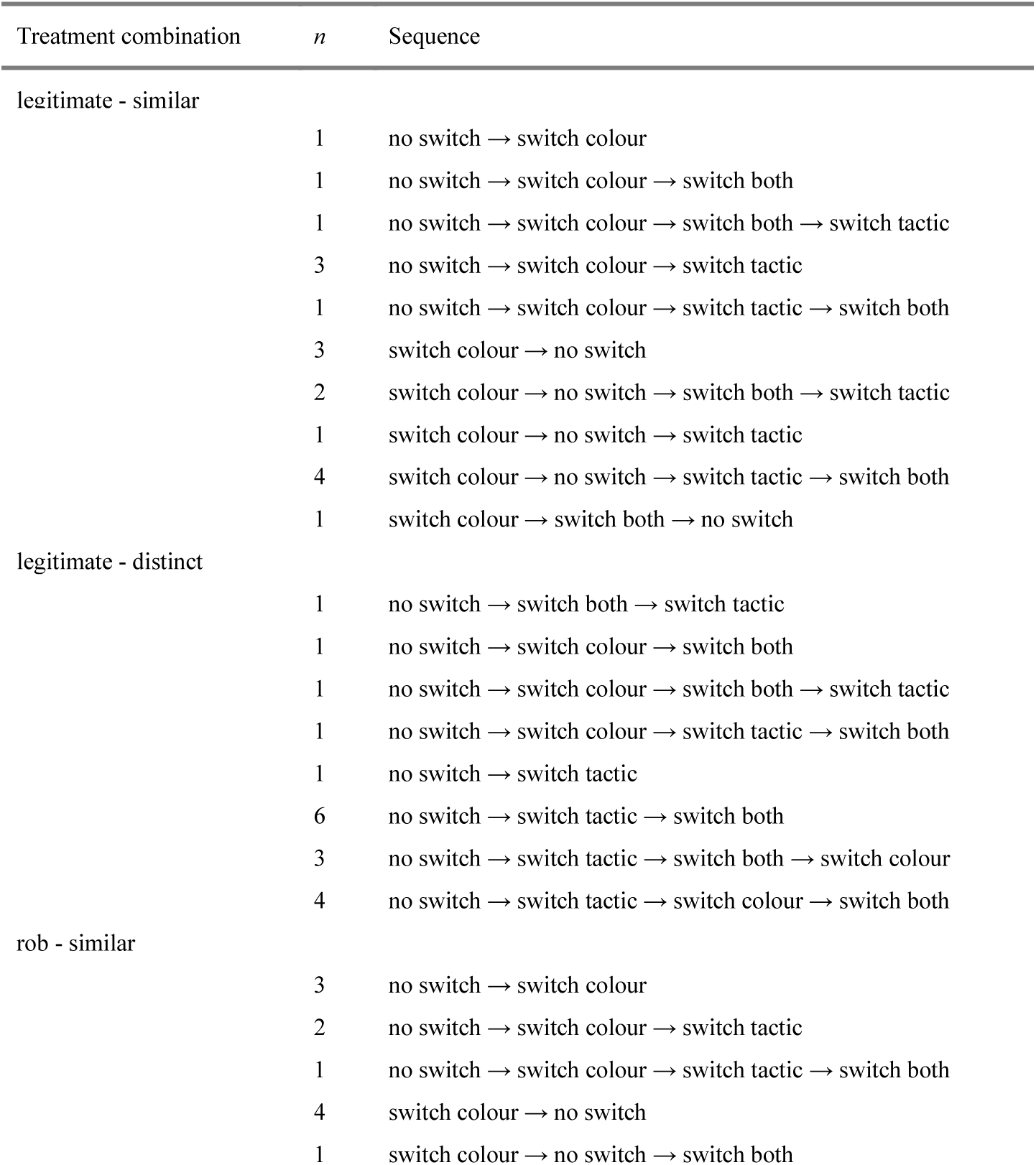
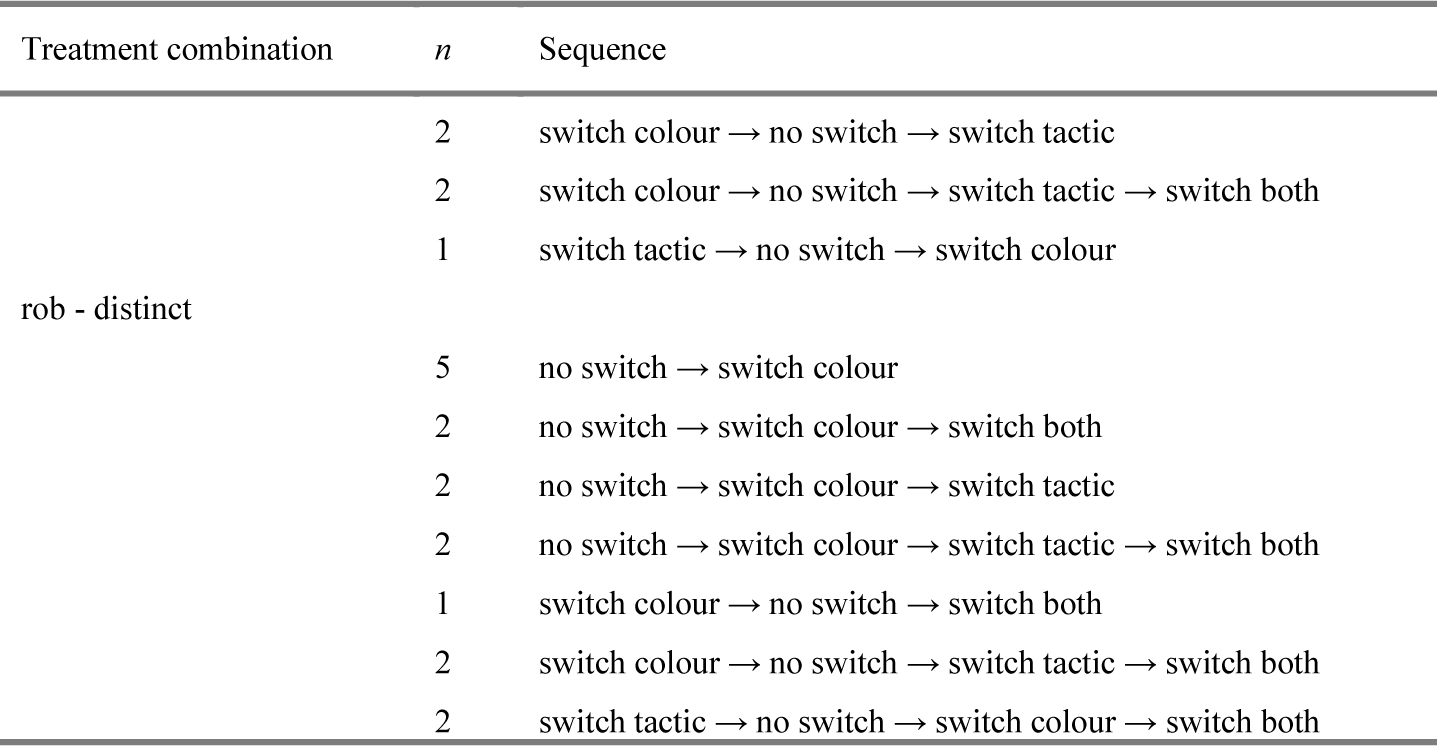
Behavioural sequences of the initial occurrences of each attempt type within different treatment combinations and its sample sizes.

**Table A4.** Pairwise comparisons with Tukey HSD adjustment on the proportion of “switch both flower colour and handling tactic” among treatment combinations.

| Pairwise comparison | Estimate | SE | *z* | *P* |
| --- | --- | --- | --- | --- |
| **legit-similar vs. legit-distinct** | **-1.501** | **0.357** | **-4.198** | **0.0002** |
| legit-similar vs. rob-similar | 0.892 | 0.581 | 1.536 | 0.4159 |
| legit-similar vs. rob-distinct | -0.040 | 0.445 | -0.090 | 0.9997 |
| **legit-distinct vs. rob-similar** | **2.393** | **0.530** | **4.516** | **<0.0001** |
| **legit-distinct vs. rob-distinct** | **1.461** | **0.376** | **3.888** | **0.0006** |
| rob-similar vs. rob-distinct | -0.933 | 0.592 | -1.577 | 0.3921 |

Significant *P*-values at 0.05 level are highlighted in bold. Estimates are given in a log-odd ratio.

**Figure A3.**
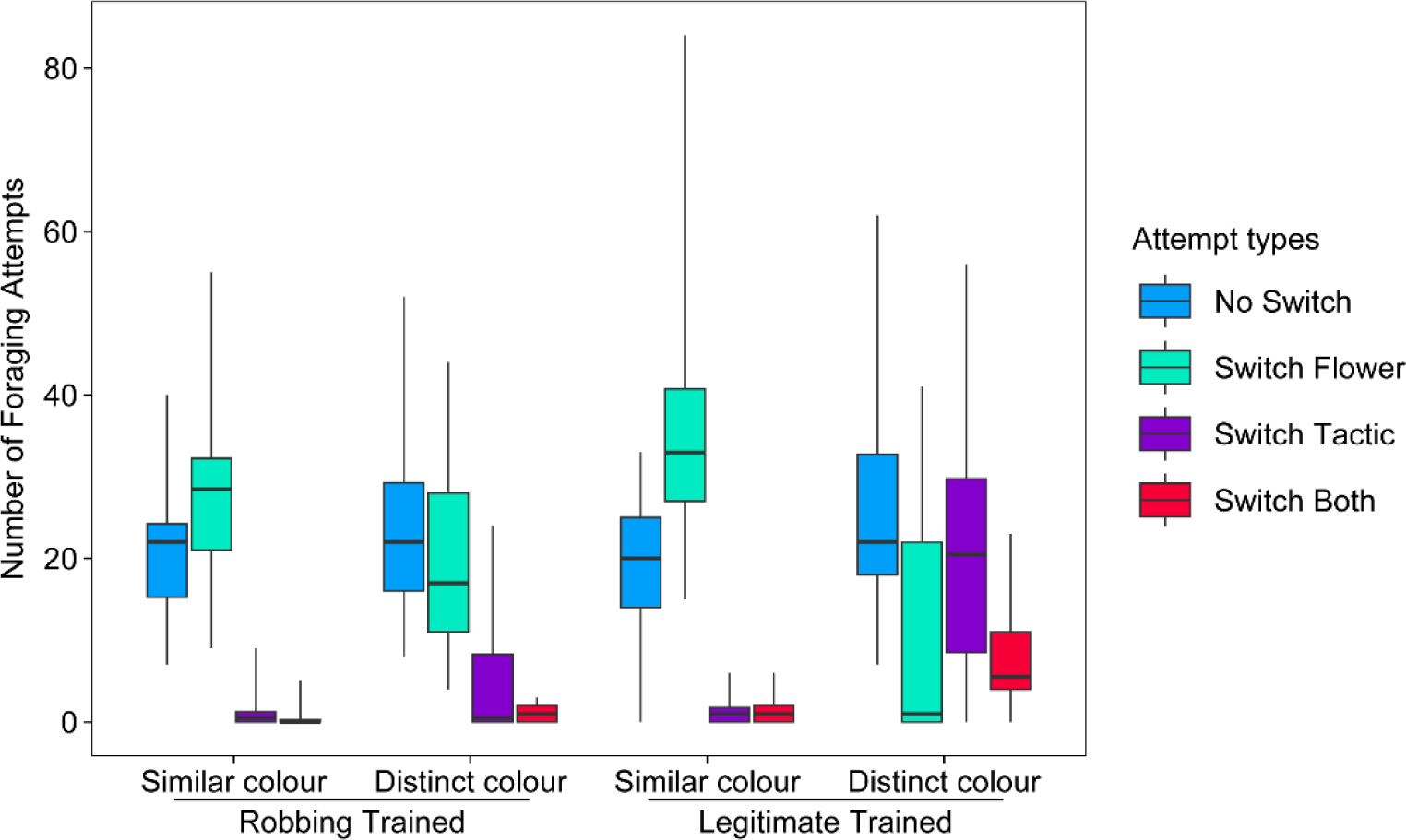
Distribution of attempt types by tactic-colour treatment combinations. Boxes indicate medians and 25% and 75% percentiles of the mean number of the four attempt types in each of the four combinations. Whiskers indicate the maximal and minimal data ranges.

## References

Baek, M., Bish, S. E., Giebink, N. W., & Papaj, D. R. (2023). The interplay of experience and pre-existing bias in nectar-robbing behavior by the common eastern bumble bee. Behavioral Ecology and Sociobiology, 77(3), 37. 10.1007/s00265-023-03313-x

Balfour, N. J., Gandy, S., & Ratnieks, F. L. W. (2015). Exploitative competition alters bee foraging and flower choice. Behavioral Ecology and Sociobiology, 69(10), 1731–1738. 10.1007/s00265-015-1985-y

Bronstein, J. L., Barker, J. L., Lichtenberg, E. M., Richardson, L. L., & Irwin, R. E. (2017). The behavioral ecology of nectar robbing: why be tactic constant? Current Opinion in Insect Science, 21, 14–18. 10.1016/j.cois.2017.05.013

Brooks, M. E., Kristensen, K., van Benthem, K. J., Magnusson, A., Berg, C. W., Nielsen, A., Skaug, H. J., Maechler, M., & Bolker, B. M. (2017). glmmTMB balances speed and flexibility among packages for zero-inflated generalized linear mixed modeling. The R Journal, 9(2), 378–400. 10.32614/RJ-2017-066

Cane, J. H., & Payne, J. A. (1993). Regional, annual, and seasonal variation in pollinator guilds: Intrinsic traits of bees (Hymenoptera: Apoidea) underlie their patterns of abundance at *Vaccinium ashei* (Ericaceae). Annals of the Entomological Society of America, 86(5), 577–588. 10.1093/aesa/86.5.577

Chittka, L. (1992). The colour hexagon: a chromaticity diagram based on photoreceptor excitations as a generalized representation of colour opponency. Journal of Comparative Physiology. A, Neuroethology, Sensory, Neural, and Behavioral Physiology, 170(5). 10.1007/bf00199331

Chittka, L. (2022). The Mind of a Bee. Princeton University Press.

Chittka, L., Beier, W., Hertel, H., Steinmann, E., & Menzel, R. (1992). Opponent colour coding is a universal strategy to evaluate the photoreceptor inputs in Hymenoptera. Journal of Comparative Physiology. A, Sensory, Neural, and Behavioral Physiology, 170(5), 545–563. 10.1007/BF00199332

Chittka, L., Gumbert, A., & Kunze, J. (1997). Foraging dynamics of bumble bees: correlates of movements within and between plant species. Behavioral Ecology, 8(3), 239–249. 10.1093/beheco/8.3.239

Chittka, L., Thomson, J. D., & Waser, N. M. (1999). Flower constancy, insect psychology, and plant evolution. Naturwissenschaften, 86(8), 361–377. 10.1007/s001140050636

Cnaani, J., Thomson, J. D., & Papaj, D. R. (2006). Flower choice and learning in foraging bumblebees: Effects of variation in nectar volume and concentration. Ethology, 112(3), 278–285. 10.1111/j.1439-0310.2006.01174.x

Coffey, M. F., & Breen, J. (1997). Seasonal variation in pollen and nectar sources of honey bees in Ireland. Journal of Apicultural Research, 36(2), 63–76. 10.1080/00218839.1997.11100932

Cotton, P. A. (2007). Seasonal resource tracking by Amazonian hummingbirds. The Ibis. 10.1111/j.1474-919X.2006.00619.x

Damas-Moreira, I., Riley, J. L., Carretero, M. A., Harris, D. J., & Whiting, M. J. (2020). Getting ahead: exploitative competition by an invasive lizard. Behavioral Ecology and Sociobiology, 74(10). 10.1007/s00265-020-02893-2

Darst, C. R., & Cummings, M. E. (2006). Predator learning favours mimicry of a less-toxic model in poison frogs. Nature, 440(7081), 208–211. 10.1038/nature04297

de Premorel, G., Giurfa, M., Andraud, C., & Gomez, D. (2017). Higher iridescent-to-pigment optical effect in flowers facilitates learning, memory and generalization in foraging bumblebees. Proceedings of the Royal Society B: Biological Sciences, 284(1865). 10.1098/rspb.2017.1097

Dedej, S., & Delaplane, K. S. (2005). Net energetic advantage drives honey bees (*Apis mellifera* L) to nectar larceny in *Vaccinium ashei* Reade. Behavioral Ecology and Sociobiology, 57(4), 398–403. 10.1007/s00265-004-0852-z

Dornhaus, A., & Chittka, L. (1999). Evolutionary origins of bee dances. Nature, 401(6748), 38–38. 10.1038/43372

Dukas, R., & Waser, N. M. (1994). Categorization of food types enhances foraging performance of bumblebees. Animal Behaviour, 48(5), 1001–1006. 10.1006/anbe.1994.1332

Dyer, A. G., & Chittka, L. (2004). Fine colour discrimination requires differential conditioning in bumblebees. Naturwissenschaften, 91(5), 224–227. 10.1007/s00114-004-0508-x

Forster, C. Y., Middleton, E. J. T., Gloag, R., Hochuli, D. F., White, T. E., & Latty, T. (2023). Impact of empty flowers on foraging choice and movement within floral patches by the honey bee, *Apis mellifera*. Insectes Sociaux. 10.1007/s00040-023-00934-3

Fox, J., & Weisberg, S. (2018). *An R Companion to Applied Regression* (3rd ed). SAGE Publications. https://play.google.com/store/books/details?id=uPNrDwAAQBAJ

Ghirlanda, S., & Enquist, M. (2003). A century of generalization. Animal Behaviour, 66(1), 15–36. 10.1006/anbe.2003.2174

Giurfa, M., Núñez, J., Chittka, L., & Menzel, R. (1995). Colour preferences of flower-naive honeybees. Journal of Comparative Physiology. A, Neuroethology, Sensory, Neural, and Behavioral Physiology, 177(3), 247–259. 10.1007/BF00192415

Grüter, C., & Ratnieks, F. L. W. (2011). Flower constancy in insect pollinators: Adaptive foraging behaviour or cognitive limitation? Communicative & Integrative Biology, 4(6), 633–636. 10.4161/cib.16972

Gumbert, A. (2000). Color choices by bumble bees (*Bombus terrestris*): innate preferences and generalization after learning. Behavioral Ecology and Sociobiology, 48(1), 36–43. 10.1007/s002650000213

Heinrich, B. (1976). The foraging specializations of individual bumblebees. Ecological Monographs, 46(2), 105–128. 10.2307/1942246

Heinrich, B. (1979). “majoring” and “minoring” by foraging bumblebees, *Bombus vagans*: An experimental analysis. Ecology, 60(2), 245–255. 10.2307/1937652

Howard, S. R., Shrestha, M., Schramme, J., Garcia, J. E., Avarguès-Weber, A., Greentree, A. D., & Dyer, A. G. (2019). Honeybees prefer novel insect-pollinated flower shapes over bird-pollinated flower shapes. Current Zoology, 65(4), 457–465. 10.1093/cz/zoy095

Inouye, D. W. (1980). The terminology of floral larceny. Ecology, 61(5), 1251–1253. 10.2307/1936841

Irwin, R. E., Brody, A. K., & Waser, N. M. (2001). The impact of floral larceny on individuals, populations, and communities. Oecologia, 129(2), 161–168. 10.1007/s004420100739

Irwin, R. E., Bronstein, J. L., Manson, J. S., & Richardson, L. (2010). Nectar robbing: ecological and evolutionary perspectives. Annual Review of Ecology, Evolution, and Systematics, 41, 271–292. 10.1146/annurev.ecolsys.110308.120330

Ishii, H. S., & Kadoya, E. Z. (2016). Legitimate visitors and nectar robbers on *Trifolium pratense* showed contrasting flower fidelity versus co-flowering plant species: could motor learning be a major determinant of flower constancy by bumble bees? Behavioral Ecology and Sociobiology, 70(3), 377–386. 10.1007/s00265-016-2057-7

Kawaguchi, L. G., Ohashi, K., & Toquenaga, Y. (2007). Contrasting responses of bumble bees to feeding conspecifics on their familiar and unfamiliar flowers. Proceedings of the Royal Society B: Biological Sciences, 274(1626), 2661–2667. 10.1098/rspb.2007.0860

Krishna, S., & Keasar, T. (2019). Bumblebees forage on flowers of increasingly complex morphologies despite low success. Animal Behaviour, 155, 119–130. 10.1016/j.anbehav.2019.06.028

Krishna, S., & Keasar, T. (2021). Generalization of Foraging Experience Biases Bees Toward Flowers With Complex Morphologies. Frontiers in Ecology and Evolution, 9. 10.3389/fevo.2021.655086

Kuusela, E., & Lämsä, J. (2016). A low-cost, computer-controlled robotic flower system for behavioral experiments. Ecology and Evolution, 6(8), 2594–2600. 10.1002/ece3.2062

Laverty, T. M. (1994). Bumble bee learning and flower morphology. Animal Behaviour, 47(3), 531–545. 10.1006/anbe.1994.1077

Laverty, T. M., & Plowright, R. C. (1988). Flower handling by bumblebees: a comparison of specialists and generalists. Animal Behaviour, 36(3), 733–740. 10.1016/S0003-3472(88)80156-8

Leadbeater, E., & Chittka, L. (2008). Social transmission of nectar-robbing behaviour in bumble-bees. Proceedings of the Royal Society B: Biological Sciences, 275(1643), 1669–1674. 10.1098/rspb.2008.0270

Lehrer, M., & Campan, R. (2005). Generalization of convex shapes by bees: what are shapes made of? The Journal of Experimental Biology, 208(Pt 17), 3233–3247. 10.1242/jeb.01790

Lenth, R. V. (2022). emmeans: estimated marginal means, aka least-squares means. https://CRAN.R-project.org/package=emmeans

Leonard, A. S., Dornhaus, A., & Papaj, D. R. (2011). Flowers help bees cope with uncertainty: signal detection and the function of floral complexity. The Journal of Experimental Biology, 214(Pt 1), 113–121. 10.1242/jeb.047407

Lichtenberg, E. M., Irwin, R. E., & Bronstein, J. L. (2018). Costs and benefits of alternative food handling tactics help explain facultative exploitation of pollination mutualisms. Ecology, 99(8), 1815–1824. 10.1002/ecy.2395

Lichtenberg, E. M., Irwin, R. E., & Bronstein, J. L. (2020). Bumble bees are constant to nectar-robbing behaviour despite low switching costs. Animal Behaviour, 170, 177–188. 10.1016/j.anbehav.2020.09.008

Loukola, O. J., Solvi, C., Coscos, L., & Chittka, L. (2017). Bumblebees show cognitive flexibility by improving on an observed complex behavior. Science, 355(6327), 833–836. 10.1126/science.aag2360

Lowe, A., Jones, L., Brennan, G., Creer, S., Christie, L., & de Vere, N. (2022). Temporal change in floral availability leads to periods of resource limitation and affects diet specificity in a generalist pollinator. Molecular Ecology. 10.1111/mec.16719

Lunau, K. (1992). Innate recognition of flowers by bumble bees: orientation of antennae to visual stamen signals. Canadian Journal of Zoology, 70(11), 2139–2144. 10.1139/z92-288

Lunau, K., Wacht, S., & Chittka, L. (1996). Colour choices of naive bumble bees and their implications for colour perception. Journal of Comparative Physiology A, 178(4), 477–489. 10.1007/BF00190178

Lynn, S. K., Cnaani, J., & Papaj, D. R. (2005). Peak shift discrimination learning as a mechanism of signal evolution. Evolution, 59(6), 1300–1305. 10.1111/j.0014-3820.2005.tb01780.x

Maharaj, G., Horack, P., Yoder, M., & Dunlap, A. S. (2018). Influence of preexisting preference for color on sampling and tracking behavior in bumble bees. Behavioral Ecology, 30(1), 150–158. 10.1093/beheco/ary140

Maia, R., Eliason, C. M., & Bitton, P. P. (2013). pavo: an R package for the analysis, visualization and organization of spectral data. Methods in Ecology and Evolution. 10.1111/2041-210X.12069

Mitchell, W. A. (1990). An optimal control theory of diet selection: The effects of resource depletion and exploitative competition. Oikos, 58(1), 16–24. 10.2307/3565356

Murphy, D. J., & Kelly, D. (2003). Seasonal variation in the honeydew, invertebrate, fruit and nectar resource for bellbirds in a New Zealand mountain beech forest. New Zealand Journal of Ecology, 27(1), 11–23. http://www.jstor.org/stable/24058156

Muth, F., Papaj, D. R., & Leonard, A. S. (2015). Colour learning when foraging for nectar and pollen: bees learn two colours at once. Biology Letters, 11(9), 20150628. 10.1098/rsbl.2015.0628

Ogilvie, J. E., & Forrest, J. R. (2017). Interactions between bee foraging and floral resource phenology shape bee populations and communities. Current Opinion in Insect Science, 21, 75–82. 10.1016/j.cois.2017.05.015

Ogilvie, J. E., & Thomson, J. D. (2016). Site fidelity by bees drives pollination facilitation in sequentially blooming plant species. Ecology, 97(6), 1442–1451. 10.1890/15-0903.1

Page, M. L., & Williams, N. M. (2023). Evidence of exploitative competition between honey bees and native bees in two California landscapes. The Journal of Animal Ecology, 92(9), 1802–1814. 10.1111/1365-2656.13973

Petanidou, T., Kallimanis, A. S., Tzanopoulos, J., Sgardelis, S. P., & Pantis, J. D. (2008). Long-term observation of a pollination network: fluctuation in species and interactions, relative invariance of network structure and implications for estimates of specialization. Ecology Letters, 11(6), 564–575. 10.1111/j.1461-0248.2008.01170.x

Pietrantuono, A. L., Requier, F., Fernández-Arhex, V., Winter, J., Huerta, G., & Guerrieri, F. (2019). Honeybees generalize among pollen scents from plants flowering in the same seasonal period. The Journal of Experimental Biology, 222(21). 10.1242/jeb.201335

Pleasants, J. M. (1981). Bumblebee response to variation in nectar availability. Ecology, 62(6), 1648–1661. 10.2307/1941519

Pleasants, J. M., & Zimmerman, M. (1979). Patchiness in the dispersion of nectar resources: Evidence for hot and cold spots. Oecologia, 41(3), 283–288. 10.1007/BF00377432

R Core Team. (2023). *R: A language and environment for statistical computing*. R Foundation for Statistical Computing. https://www.R-project.org/

Raine, N. E., & Chittka, L. (2007a). Flower constancy and memory dynamics in bumblebees (Hymenoptera: Apidae: Bombus). Entomologia Generalis, 29, 179–199. 10.1127/ENTOM.GEN/29/2007/179

Raine, N. E., & Chittka, L. (2007b). Pollen foraging: learning a complex motor skill by bumblebees (*Bombus terrestris*). Naturwissenschaften, 94(6), 459–464. 10.1007/s00114-006-0184-0

Rands, S. A., Whitney, H. M., & Hempel de Ibarra, N. (2023). Multimodal floral recognition by bumblebees. Current Opinion in Insect Science, 59, 101086. 10.1016/j.cois.2023.101086

Real, L., & Rathcke, B. J. (1988). Patterns of individual variability in floral resources. Ecology, 69(3), 728–735. 10.2307/1941021

Richman, S. K., Barker, J. L., Baek, M., Papaj, D. R., Irwin, R. E., & Bronstein, J. L. (2021). The sensory and cognitive ecology of nectar robbing. Frontiers in Ecology and Evolution, 9. 10.3389/fevo.2021.698137

Rodríguez, I., Gumbert, A., Hempel de Ibarra, N., Kunze, J., & Giurfa, M. (2004). Symmetry is in the eye of the beeholder: innate preference for bilateral symmetry in flower-naïve bumblebees. Naturwissenschaften, 91(8), 374–377. 10.1007/s00114-004-0537-5

Rojas-Nossa, S. V., Sánchez, J. M., & Navarro, L. (2016). Nectar robbing: a common phenomenon mainly determined by accessibility constraints, nectar volume and density of energy rewards. Oikos, 125(7), 1044–1055. 10.1111/oik.02685

Russell, A. L., Leonard, A. S., Gillette, H. D., & Papaj, D. R. (2016). Concealed floral rewards and the role of experience in floral sonication by bees. Animal Behaviour, 120, 83–91. 10.1016/j.anbehav.2016.07.024

Russo, L., Debarros, N., Yang, S., Shea, K., & Mortensen, D. (2013). Supporting crop pollinators with floral resources: network-based phenological matching. Ecology and Evolution, 3(9), 3125–3140. 10.1002/ece3.703

Rust, R. W. (1979). Pollination of Impatiens capensis: Pollinators and Nectar Robbers. Journal of the Kansas Entomological Society, 52(2), 297–308. http://www.jstor.org/stable/25083907

Schiestl, F. P., & Johnson, S. D. (2013). Pollinator-mediated evolution of floral signals. Trends in Ecology & Evolution, 28(5), 307–315. 10.1016/j.tree.2013.01.019

Simpson, B. B., & Neff, J. L. (1981). Floral rewards: alternatives to pollen and nectar. Annals of the Missouri Botanical Garden. Missouri Botanical Garden, 68(2), 301–322. 10.2307/2398800

Skorupski, P., & Chittka, L. (2010). Photoreceptor spectral sensitivity in the bumblebee, *Bombus impatiens* (Hymenoptera: Apidae). PloS One, 5(8), e12049. 10.1371/journal.pone.0012049

Stephens, R. B., Hobbie, E. A., Lee, T. D., & Rowe, R. J. (2019). Pulsed resource availability changes dietary niche breadth and partitioning between generalist rodent consumers. Ecology and Evolution, 9(18), 10681–10693. 10.1002/ece3.5587

Thomson, J. D., McKenna, M. A., & Cruzan, M. B. (1989). Temporal patterns of nectar and pollen production in *Aralia hispida*: Implications for reproductive success. Ecology, 70(4), 1061–1068. 10.2307/1941375

Valls-Fox, H., De Garine-Wichatitsky, M., Fritz, H., & Chamaillé-Jammes, S. (2018). Resource depletion versus landscape complementation: habitat selection by a multiple central place forager. Landscape Ecology, 33(1), 127–140. 10.1007/s10980-017-0588-6

van der Kooi, C. J., Dyer, A. G., Kevan, P. G., & Lunau, K. (2019). Functional significance of the optical properties of flowers for visual signalling. Annals of Botany, 123(2), 263–276. 10.1093/aob/mcy119

van der Kooi, C. J., Vallejo-Marín, M., & Leonhardt, S. D. (2021). Mutualisms and (a)symmetry in plant-pollinator interactions. Current Biology, 31(2), R91–R99. 10.1016/j.cub.2020.11.020

Waser, N. M. (1986). Flower constancy: definition, cause, and measurement. The American Naturalist, 127(5), 593–603. 10.1086/284507

